# Natural SARS-CoV-2 infections, including virus isolation, among serially tested cats and dogs in households with confirmed human COVID-19 cases in Texas, USA

**DOI:** 10.1101/2020.12.08.416339

**Authors:** Sarah A. Hamer, Alex Pauvolid-Corrêa, Italo B. Zecca, Edward Davila, Lisa D. Auckland, Christopher M. Roundy, Wendy Tang, Mia Torchetti, Mary Lea Killian, Melinda Jenkins-Moore, Katie Mozingo, Yao Akpalu, Ria R. Ghai, Jessica R. Spengler, Casey Barton Behravesh, Rebecca S. B. Fischer, Gabriel L. Hamer

**Author notes:** **One Sentence Summary:** In over 25% of homes with infected humans, pet dogs or cats had confirmed SARS-CoV-2 infections, including one cat where virus was successfully isolated for the first time from a companion animal.

## Abstract

The natural infections and epidemiological roles of household pets in SARS-CoV-2 transmission are not understood. We conducted a longitudinal study of dogs and cats living with at least one SARS-CoV-2 infected human in Texas and found 47.1% of 17 cats and 15.3% of 59 dogs from 25.6% of 39 households were positive for SARS-CoV-2 via RT-PCR and genome sequencing or neutralizing antibodies. Virus was isolated from one cat. The majority (82.4%) of infected pets were asymptomatic. Re-sampling of one infected cat showed persistence of viral RNA at least 32 d-post human diagnosis (25 d-post initial test). Across 15 antibody-positive animals, titers increased (33.3%), decreased (33.3%) or were stable (33.3%) over time. A One Health approach is informative for prevention and control of SARS-CoV-2 transmission.

## Main Text

The global pandemic of coronavirus disease 2019 (COVID-19), caused by the novel severe acute respiratory syndrome virus 2 (SARS-CoV-2), is an unprecedented challenge to health, the economy, and nearly all aspects of society. Phylogenetic studies that reconstruct the evolutionary relationships between SARS-CoV-2 and its closest relatives suggest the reservoir of SARS-CoV-2 was an animal host, likely horseshoe bats (*Rhinolophus* spp.), although the virus may have emerged in the human population by way of a yet unidentified intermediate host (*1*). As the global pandemic continues, studies have established that this virus is zoonotic, with experimental or epidemiological studies discovering that several mammalian groups, including primates, felids, mustelids, and some species of rodent, lagomorph, and bat are susceptible to natural or experimental infection (*2*). Across the globe, there are two common groups where suspected human-to-animal transmission events have been repeatedly detected: farmed mink and companion animals. Outbreaks have occurred on mink farms in the Netherlands, Denmark, Spain, Italy, Sweden, Greece, and the United States between April and November 2020. Outbreaks in Europe led to widespread culling and moratoria being placed on the mink industry (*3*). With respect to companion animals, over 10 countries to-date have documented natural infections of dogs and cats, often associated with exposure to a person with COVID-19 (*4*). For example, the first cases of companion animal infection with SARS-CoV-2 in the United States were two cats in New York that developed respiratory signs after exposure to their owners with suspected or confirmed COVID-19 (*5*).

Experimentally, cats have been shown to be highly susceptible to infection with SARS-CoV-2, and can transmit the virus to other cats under laboratory conditions via both direct and indirect contact (*6–9*). Bosco-Lauth et al. (*7*) demonstrated that neutralizing antibodies protected cats against subsequent challenge. There is also evidence from laboratory challenge studies that dogs have lower susceptibility with limited viral replication, although seroconversion with neutralizing antibodies has been confirmed for both dogs and cats (*6, 7*).

Despite the potential public health and animal health implications of SARS-CoV-2 infections in companion animals, few epidemiological investigations of companion animals living in household environments are available. Most evidence of natural animal exposures and infections comes from opportunistic case studies (*10*), testing of animals presenting for routine veterinary visits, or studies in congregate animal settings (*11*). Focused studies of animals with known exposure to people with COVID-19 have the potential to quantify probability of infection in animals that have sustained contact with an infected person, and therefore are critical for understanding the potential for companion animals to serve as reservoirs for the virus. Furthermore, there are no published reports of SARS-CoV-2 respiratory infections or viral isolation from companion animals in household transmission studies in the United States. The objective of this study was to establish an epidemiological investigation program based on the active surveillance of dogs and cats from households with SARS-CoV-2-infected owners in order to quantify the prevalence of domestic animal infection in these high risk natural environments in Texas, a state with high numbers of reported human cases in summer 2020 (*12*).

## Results

### Demographic data

Between June 24–July 31, 2020, 76 pets from 39 households were sampled in Brazos County, Texas. Diverse breeds of dogs and cats were represented; cats ranged in age from 3 months to 10 years, and dogs ranged in age from 1.5 months to 18 years. Specimens were collected from pets 3-27 days after the human household member received positive test result (mean 9 days; median 8 days), with up to three follow-up specimen collections for some pets occurring through September 7, 2020.

### Infection prevalence and viral isolation

SARS-CoV-2 infection was confirmed by PCR and sequencing in 3 of 17 (17.6%) cats and 1 of 59 (1.7%) dogs sampled from 4 of 39 (10.3%) separate households; these four animals met the United States Department of Agriculture (USDA) animal case definition for SARS-CoV-2 infection, with positive respiratory and/or rectal swabs (Table 1). An additional two dogs that lived together tested positive by rectal and body swabs (TAMU-044) or body swab only (TAMU-043), but did not meet case definition (e.g. sequence data) likely due to presence of low concentration of viral targets based upon high Ct values. Further, these two dogs had no evidence of neutralizing antibodies and did not develop any upon re-sampling, supporting the absence of prior infection and that detection by PCR likely reflects living in a contaminated environment.

**Table 1.**
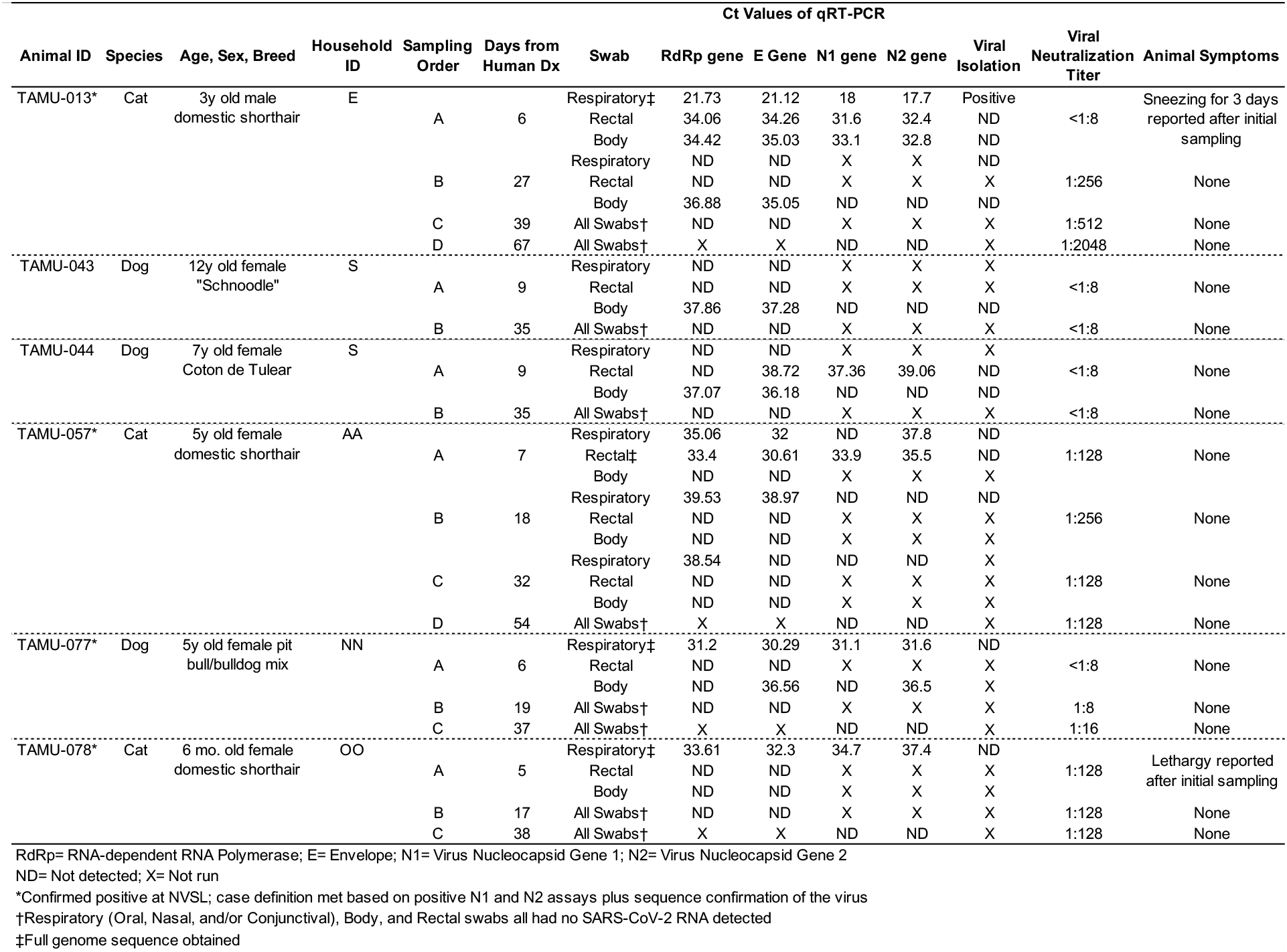
Pets with positive screening or confirmatory qRT-PCR results for detection of SARS-CoV-2 virus from households with at least one confirmed case of human COVID-19 in Brazos County, Texas, at first (A) and subsequent (B–D) specimen collection visits from June-September, 2020.

In one (cat TAMU-013) of four SARS-CoV-2 confirmed animals based on sequence data, all 3 swab types (respiratory, rectal, and body) tested positive by all PCR tests. Additionally, this same cat was the only animal to test positive in the pan-coronavirus conventional PCR, resulting in an RdRp gene partial sequence with 100% sequence homology to several SARS-CoV-2 isolates from human cases in the United States (Genbank accession no. MT911466). Of the three other confirmed cases, all had positive respiratory swabs, with one cat also testing positive by PCR via rectal swab, and a dog from which body swabs tested positive by PCR (Table 1). All four confirmed cases were sampled within 7 days of the reported diagnosis of their owner (Table 1). Of the four households with a confirmed case, the median number of days between a positive human test result and the sampling of pets was 6 days; in contrast, of the 35 households with no pet meeting case definition based on PCR, the interval was 8 days (Mann-Whitney U test, P=0.08).

Virus isolation was attempted from nine specimens (Table 1), including six specimens from the four cases confirmed by sequence, plus three additional specimens positive by initial PCR only. SARS-CoV-2 was isolated only from the respiratory swab of TAMU-013 on second passage with no cytopathic effects (CPE) noted on day 7. Virus recovery was confirmed by PCR (Ct value reduction from original inoculum to cell suspension of second passage).

### Sequence analysis

Coding-complete genome sequence was obtained for all four confirmed positive animals from respiratory or rectal swabs collected at the first visit (Table 1). All genomes shared high levels of identity to SARS-CoV-2 (99.97-99.98%) and contained 12-16 mutations compared to reference sequence Wuhan-Hu-1. The animal viruses belong to clades G (n=1), GH (n=2), and GR (n=1) which correspond to the predominant clades observed for human samples from Texas during the same time period (Figure 2). No mutations were observed in the spike protein receptor binding domains.

### Virus neutralization and household patterns

Viral neutralization was attempted for 75 of the 76 animals in the study; one cat was fractious at the time of sampling, so no blood was obtained. At baseline specimen collection, 14 of 75 (18.7%) pets had SARS-CoV-2 neutralizing antibodies, including 7 of 16 (43.8%) cats and 7 of 59 (11.9%) dogs from 10 of 39 (25.6%) households. These included 2 of the 4 (50%) PCR-confirmed positive case animals (Table 2). Virus neutralization titers ranged from 1:16 to 1:128 upon initial sampling.

**Table 2.**
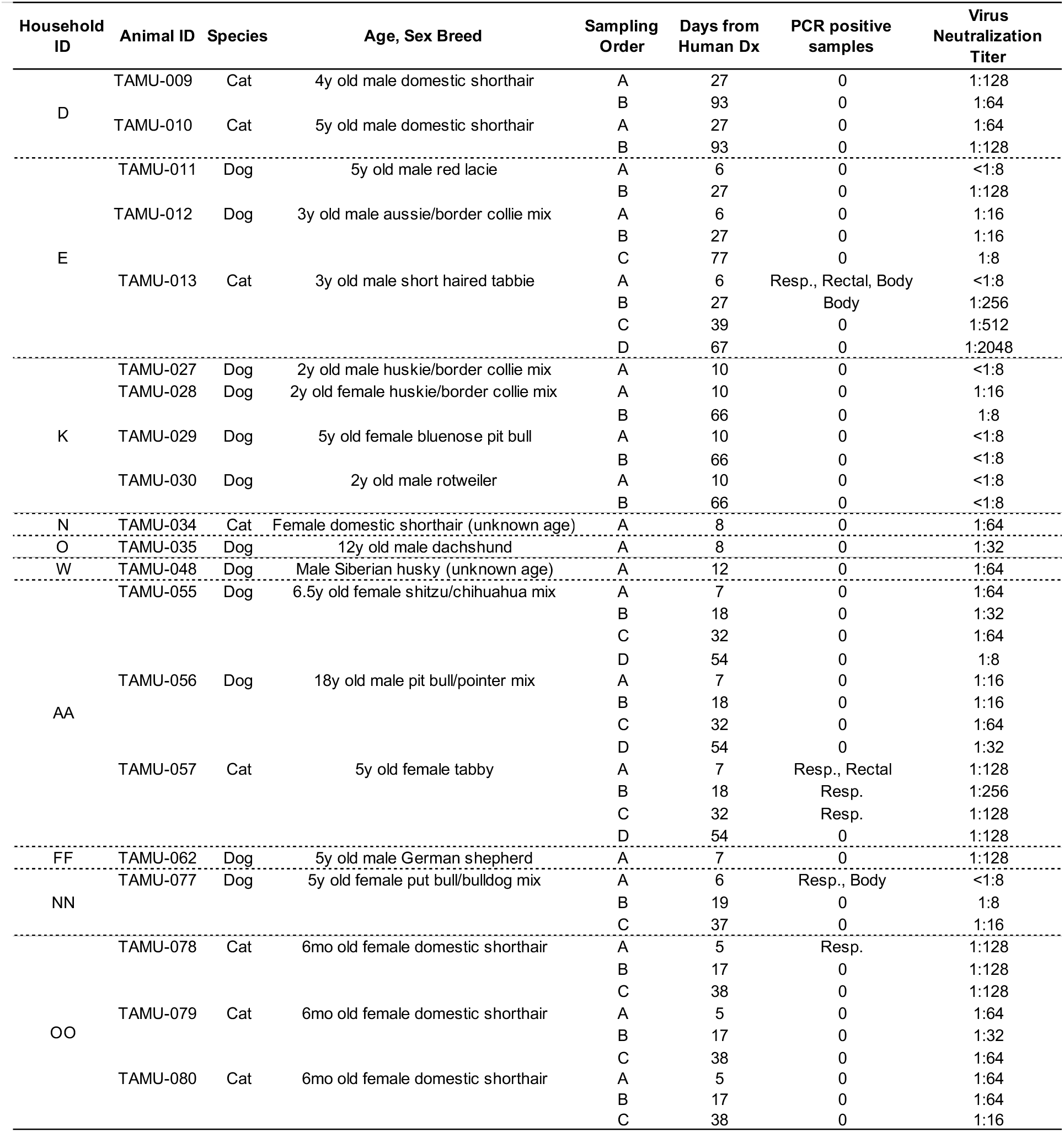
Pets with SARS-CoV-2 neutralizing antibody titers from households with at least one confirmed case of human COVID-19 in Brazos County, Texas, at first (A) and subsequent (B–D) sampling visits from June-August, 2020. All households with at least one VN-positive animal are shown, with all animals living in the household tested one or more times. For the column of PCR results, any positive PCR result in any of the screening or confirmatory assays is considered positive.

Among animals in multi-pet households (Figure 1), there are some instances where all animals were negative by PCR, yet positive by VN. For example, both cats from household D were PCR-negative yet VN-positive, and were sampled 27 days after owner diagnosis. In household K, all four dogs were PCR-negative and one was VN-positive, and were sampled 10 days after owner diagnosis. In the three multi-pet households, each had a single cat test PCR-positive at the initial visit with neutralizing antibodies found in 1 of 3 (household E; sampled 6 days after owner diagnosis) and in 3 of 3 pets at the second sampling (household AA; sampled 7 days after owner diagnosis, and household OO; sampled 5 days after owner diagnosis).

**Fig. 1.**
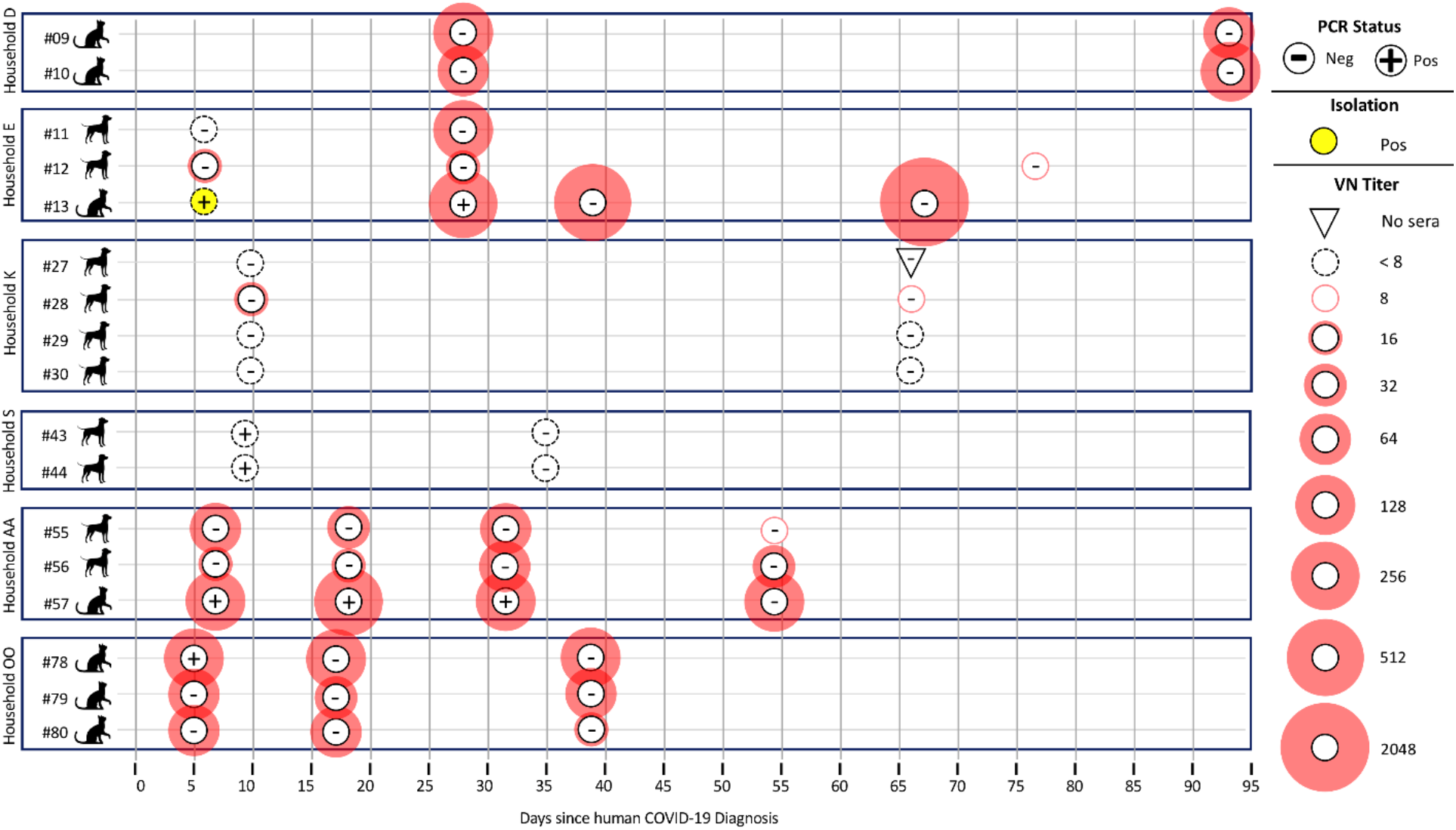
Multi-pet households in which at least one pet in the house was initially positive by PCR analyses or virus neutralization. All pets in the household were followed longitudinally to track duration of positive test results and changes in antibody titers over time. PCR status is based on preliminary screening Ct of <40 on RdRp, E, N1 and/or N2 gene target from any diagnostic swab type [respiratory (nasal/oral/conjunctival), rectal], or body swab; positive results here do not necessarily indicate USDA case definition has been met. Virus neutralization titer was determined by a two-fold serial dilution of sera.

**Fig. 2.**
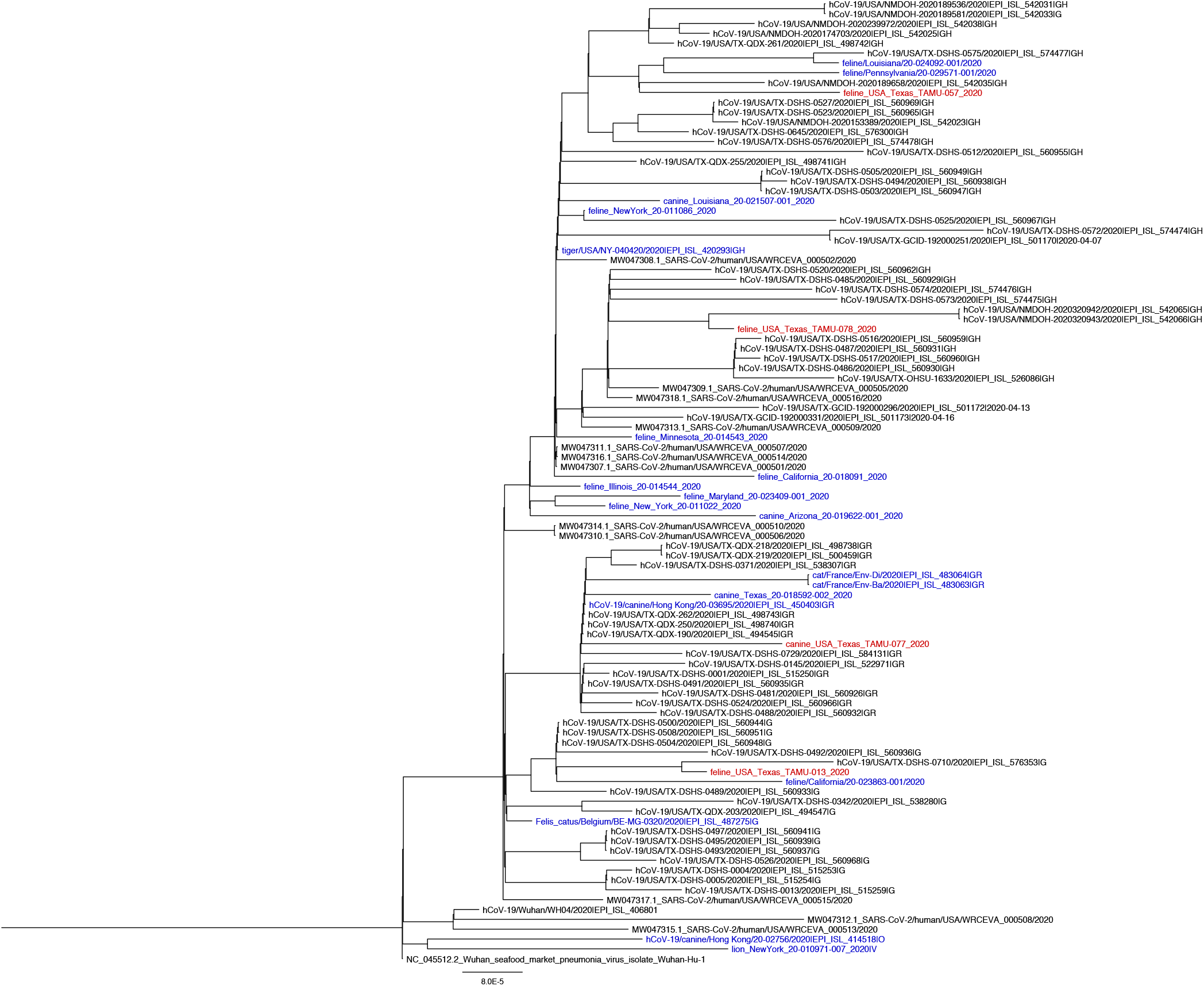
Phylogenetic analysis of SARS-CoV-2 whole genome sequences obtained from three cats and one dog from this study (red; animal ID: TAMU-013, 057, 077, 078) and SARS-CoV-2 genomes from feline and canine hosts based on prior studies (blue) and from humans from Texas (black). The analysis used RAxML with the GTRCAT model (*25*).

### Longitudinal sampling of infected animals

Animals testing positive by PCR and/or VN, and all other animals living in the household with positive animals, were re-sampled up to three times within 2 months of the first specimen collection to describe the natural history of infection in terms of viral RNA and antibody titers over time, resulting in 30 follow-up events. In 2 of the 4 animals initially confirmed by sequence, the follow-up samplings were negative by PCR (Table 1). However, for one cat (TAMU-013), the body swab was also PCR-positive (for 2 of 4 viral targets) from the second sample acquired 3 weeks later (27 days after owner’s diagnosis) with comparable Ct values. In one other cat (TAMU-057), the respiratory swab was positive by PCR (for 2 of 4 viral targets) in the second sample that was acquired 1.5 weeks later but with increased Ct values, and also at third sampling (for 1 of 4 viral targets) nearly four weeks after the first sample but with a high Ct value. This represents the longest published duration of detection of viral RNA by preliminary screening in a companion animal sample to date, with the last PCR positive result recorded 32 days following owner’s diagnosis (Table 2).

Among all 15 animals that had sequential serum samples collected, antibodies increased over time in five (33%), decreased in five (33%), and remained stable in the remaining five animals (33%; Table 2). One cat (TAMU-057), which was also PCR-positive at initial visit, had VN titers of 1:128, 1:256, 1:128, and 1:128 at four sequential visits from 7-54 days post owner diagnosis. There was also evidence of seroconversion in individual animals; for example, dog TAMU-077 was PCR-positive and VN-negative at first visit, had a VN titer of 1:8 by the second visit 13 days later, and a titer of 1:16 by the third visit 18 days later.

In examining the multi-pet houses that were re-sampled, we see that the number of VN-positive animals in the house increased over time. For example, household E, where one cat and two dogs live, had an initial sampling status of PCR-positivity in the cat, with only one dog testing VN-positive. By three weeks later, the cat was still PCR-positive and all three animals were VN-positive. Additional sampling of the cat (TAMU-013) showed seroconversion followed by increasing VN titer (negative, 1:256, 1:512, and 1:2048 at the four visits, respectively).

### Pet signs of disease

Among the four pets confirmed by sequence and two others that were presumptive positive by PCR but were not confirmed, owners reported all pets to be asymptomatic prior to the time of first sampling. Subsequently, one cat (TAMU-013) was reported to sneeze for approximately three days following the first sampling. Additionally, another cat (TAMU-078) was reported to be more sleepy than normal for 2 weeks following the first sampling. Among the additional 13 animals that were VN-positive during initial or re-sample events, a single dog (TAMU-028) was reported to sneeze prior to initial sampling. When re-sampled all animals were reported to be in good health.

## Discussion

There are few active surveillance studies assessing SARS-CoV-2 infection in companion animals in households with documented human infections. In France, one study of 9 cats and 12 dogs owned by people with COVID-19 showed no evidence of SARS-CoV-2 viral RNA or antibodies (*13*), while another study of 22 cats and 11 dogs found a single cat (4.5%) positive for SARS-CoV-2 by RT-PCR (*14*). A study in Hong Kong, China, documented 2 of 15 (13.3%) dogs living in homes with human COVID-19 had SARS-CoV-2 viral RNA in nasal, oral, and rectal swabs and also demonstrated measurable antibody titers on subsequent samples (*15*). In another Hong Kong study, 6 of 50 (12%) cats living with humans with SARS-CoV-2 infection were positive by RT-PCR (*16*).

The current study rapidly deployed a field team to assess infection among pets living with laboratory-confirmed SARS-CoV-2-infected owners and used a comprehensive sampling and testing protocol, a strength of this study for increased probability of timely SARS-CoV-2 detection. We focused sampling on dogs and cats at high risk of exposure to SARS-CoV-2 through their owners at a time where the rate of human community transmission in Texas was high [average of 27.3 cases/100,000 people in the state during the late June-July study period (*12*)]. Of 17 cats, live SARS-CoV-2 was recovered from respiratory swab from one (5.9%), 16.7% tested positive by PCR from one or more swab type, and 41.2% had neutralizing antibodies. Of 59 dogs, 1.7% tested positive by PCR from one or more swab sample type and 11.9% had neutralizing antibodies. The phylogenetic analysis from this study shows 4 of the SARS-CoV-2 genomes recovered from three cats and one dog are of unique lineages and likely represent community spill-back. In other studies assessing seroprevalence in companion animals living in households with SARS-CoV-2 infected owners, seropositivity of 4-23.5% has been reported in cats and 13-20% in dogs (*16–20*). Taken together, the current study suggests human-to-animal transmission of SARS-CoV-2 may occur more often for cats than previously recognized.

To our knowledge this study documents the first isolation of SARS-CoV-2 from an animal naturally exposed to a human with laboratory-confirmed SARS-CoV-2 infection living in the same household. Virus was successfully isolated from the swab sample with the lowest Ct value observed in this study, consistent with what has been reported on virus isolation from human-derived specimens, which has been successful from samples with Ct values under 25 (*21*) or 10^6^ RNA copies per mL of sample (*22*). Despite recovery of live SARS-CoV-2 from one feline respiratory sample, this study provides no evidence that companion animals play a role in spreading the virus to humans or to other animals. Three out of 4 households with confirmed cases were multi-pet households, each with two other animals in the household. All pets in each of these households had SARS-CoV-2 neutralizing antibodies from follow-up serum collections. We did not evaluate the chain or timing of infection and cannot speculate whether human-to-animal or animal-to-animal transmission occurred. Nevertheless, CDC recommends that people with suspected or confirmed COVID-19 isolate from their pets, just as they would from other members of their household, to reduce the potential for human-to-animal transmission (*23*).

One factor likely contributing to the proportion of household companion animals testing positive for viral RNA or infectious virus is the relatively short window for sampling following human symptoms or diagnosis. Specifically, all four animals meeting case definition by sequence were sampled within 7 days of owner diagnosis. A study in dogs in Hong Kong found similar results, with SARS-CoV-2 RNA detection in a dog sampled 14 days following initial symptoms of the human COVID-19 patient and one day following confirmed diagnosis. Experimental infection studies indicate the window of SARS-CoV-2 shedding in cats is up to 7 days (*7*). Another factor contributing to the detection of positive animals in the current study is the multiple swab types (3 to 5) taken from each animal and the two gene targets used for screening all animals and the two additional gene targets used to confirm presumptive positives. With this approach we detected viral RNA by at least one viral PCR target from three sequential sampling events spaced 32 days apart for a single cat.

Across the study, there were four detections from body (fur) swabs by PCR. Two of these samples were in the same cat (TAMU-013); the first positive body swabs coincided with the initial sampling event when the cat’s respiratory and rectal swabs were also strongly positive, and the second occurred three weeks later when only the body swab was positive. Given self-licking and grooming behavior of cats, the viral RNA on the fur may reflect self-contamination or reflect living in a household environment where the virus had persisted in prior weeks. The other samples were from two dogs living in the same household (household S); one dog also had a positive rectal swab and the other tested positive only on the body swab; however, neither dog met the case definition. Upon follow-up sampling 3.5 weeks later, both dogs were PCR-negative and remained VN-negative, suggesting they were not infected and the initial detections likely reflected living in contaminated environment. Attempts to isolate virus from all four PCR-positive fur samples were unsuccessful providing further evidence that pet fur is not a fomite for SARS-CoV-2 transmission.

This study documents that over 25% of households sampled had pets with neutralizing antibodies. The neutralizing antibody results in this study were based on an inclusive cut-off of 1:8 to consider an animal positive. In general, the titers of neutralizing antibodies in animals presented here seem to fluctuate along time with two-thirds of animals presenting increase or decrease in antibody titers. More studies evaluating the dynamic of infection not only in pets, but also in their respective owners would help elucidate the factors involved in the antibody titer variation observed in companion animals. Although this neutralization test was designed to be specific to SARS-CoV-2, and no cross-reactivity between SARS-CoV-2 and type I and II of feline peritonitis virus was detected in a recent study using similar methods in a cat in Wuhan (*11*), more comprehensive serology assays including other coronaviruses to confirm monotypic reactions to SARS-CoV-2 may be warranted.

Very few case studies of natural infection in cats and dogs document severe clinical outcomes, and those that have reveal that co-morbidities likely played a contributing factor in illness or death (*15, 24*). In our study, across all 17 animals that were PCR-positive, VN-positive, or both, only three animals were reported to have mild signs of disease, including 3 days of sneezing in a PCR-positive cat; lethargy in a PCR-positive cat; and sneezing in an antibody-positive dog. Importantly, upon resample, these and all other animals in the study were reported to be in good health. Based on OIE reports, slightly less than half of all animals reported with SARS-CoV-2 infection have clinical signs, which may include fever, coughing, difficulty breathing or shortness of breath, lethargy, sneezing, nasal discharge, ocular discharge, vomiting, and diarrhea (*4*). Our results, which reveal a much higher proportion of asymptomatic animals, suggest that infected companion animals showing no clinical signs or only mild, transient illness may be more numerous than global reporting currently captures. Further active surveillance efforts would more accurately determine the true burden of this virus within the companion animal population.

Study limitations include that it is unknown how many SARS-CoV-2-positive humans lived in each household, or their duration or nature of their symptoms while interacting with pets. Further, specimens or sequences from the human SARS-CoV-2 infections in the same households were not available for alignment with the sequences we generated from the pets, which would be useful for understanding transmission. Additionally, because animals associated with initial negative results in households with other negative animals were never re-sampled, we may have missed the detection of infection if animals seroconverted at a later date.

The present study advances our understanding of the transmission risk between people and their pets during the COVID-19 pandemic. Moreover, it underscores the need for a One Health approach (*37*), both in epidemiological investigations and in prevention and control measures as well as pandemic preparedness for SARS-CoV-2 and other emerging zoonotic infectious diseases.

## Materials and Methods

### Animal recruiting and sampling

Study enrollment criteria included any dog or cat living in the same household as a person with a confirmed (PCR-positive) SARS-CoV-2 infection; no restrictions were made based on breed, age, vaccination status or medical history of animals. Individuals that tested positive for SARS-CoV-2 were contacted via phone by the Brazos County Health Department (BCHD) as part of a public health case investigation. Individuals were asked if they owned pet dogs or cats, and if they did, if they wished to learn more about enrolling their pets in a TAMU research project sampling animals for SARS-CoV-2. Interested pet owners were provided the project website (tx.ag/BCSCovidResearch), and the contact information of consenting pet owners was provided to the TAMU investigation team. Pet owners were administered a short questionnaire by phone including pet signalment (breed, age, sex), vaccination history, pet symptoms, date of positive human test result (which was cross-checked with BCHD), and were read the details of the informed consent form, after which a visit was arranged to sample the animal(s) at their household.

All samples were obtained from privately-owned animals in adherence with animal use protocols approved by the Texas A&M University’s Institutional Animal Care and Use Committee and Clinical Research Review Committee on May 14, 2020 (2018-0460 CA). Written consent was received for each pet from the owner. For sample collection at the household, three swab samples were obtained from each animal: (i) respiratory; (ii) rectal; and (iii) external body (fur). The respiratory sample was a combination of an oral/oropharyngeal swab, a nasal/nasopharyngeal swab, and a conjunctival swab (cats only). The rectal sample included a single swab inserted up to 1.5cm in the rectum and around the external surface. Due to concerns that emerged early in the pandemic of animals serving as fomites for the virus, body (fur) swabs were also collected by rubbing two swabs over the animal’s scruff, ears, neck, back, and abdomen. All swabs were 5.2 mm diameter standard polyester tipped applicators with polystyrene handles (Puritan Medical Products, Guilford, ME) except for cat and small dog respiratory samples, where 3.3 mm diameter polyester swabs with propylene handles (Constix^®^, Contec, Spartanburg, SC) were used. Swabs were submerged immediately into 3 mL of viral transport media (VTM; made following CDC SOP#: DSR-052-02) and left in media until aliquots were prepared within 24 hours. Blood was collected via cephalic, jugular, or medial saphenous venipuncture into clot activator and EDTA-tubes. All samples were kept in a cooler with ice packs until returned to the lab. Swabs were placed in a −80°C freezer. Blood samples were centrifuged and aliquots were prepared of serum, clot, whole blood, plasma, and red blood cells which were stored at −80°C.

For households with animals that tested positive (viral RNA or antibodies), efforts were made to longitudinally re-sample all animals living in the household regardless of the initial testing result of each individual animal. Repeat specimen collections followed a minimum of one week from the date of initial sample collection and occurred up to three times per household within two months of the first specimen collection event. At each follow-up visit, identical sets of specimen types were collected, and owners were asked about any changes in their pet(s) health. For some follow-up events, an updated protocol was implemented, which included collection of oral/oropharyngeal, nasal/nasopharyngeal, and conjunctival swabs separately, rather than combining these swabs into a single respiratory sample.

### Molecular testing

All specimens were evaluated for the presence of SARS-CoV-2 viral RNA at Texas A&M University using the following protocols. The VTM was homogenized and a 400 μL aliquot was removed. Viral RNA was extracted using a MagMAX CORE Nucleic Acid Purification Kit on a 96-well Kingfisher Flex System (ThermoFisher Scientific, Waltham, MA). A subset of samples was tested for RNA concentration on an Epoch Microplate Spectrophotometer (BioTek, Winooski, VT). All samples were screened on two separate qRT-PCRs targeting the RNA-dependent RNA polymerase (RdRp) gene and the E gene (*26, 27*). Briefly, a 25μL reaction included 5μL of sample RNA, 6.25μL of 4X RT-Buffer (TaqMan Fast Virus 1-Step Master Mix, ThermoFisher Scientific), 600nM of the forward primer (RdRp_SARSr-F), 800nM of the reverse primer (RdRp_SARSr-R), and 100nM of the probe (RdRp_SARSr-P2); a control plasmid containing a portion of the RdRp gene served as positive control. The E gene 25μL reaction consisted of 5μL of viral RNA, 6.25μLof 4X RT-Buffer, 400nM of the forward primer (E_Sarbeco_F), 400nM of the reverse primer (E_Sarbeco_R), and 200nM of the probe (E_Sarbeco_P1); a control plasmid containing the complete envelope gene served as positive control. All primers, probes, and positive controls were from IDT (Integrated DNA Technologies, Coralville, IA). Both reactions followed the previously published thermocycling conditions (*26, 27*) which consisted of 50°C for 30 min for reverse transcription, followed by 95°C for 15 min for RT inactivation/initial denaturation, then 45 cycles of 95°C for 15s, 58°C for 60s using a CFX96 Real-Time System (BIORAD, Hercules, CA).

All specimens that were non-negative using either protocol were submitted for confirmatory PCR testing at the USDA NVSL with joint approval from the Texas Department of State Health Services and the Texas Animal Health Commission. An 800 μL aliquot of VTM swab sample was sent to NVSL where two additional qRT-PCR protocols were followed, targeting virus nucleocapsid gene 1 (N1) and nucleocapsid gene 2 (N2) for specific detection of SARS-CoV-2 following the CDC’s protocol (*28*). Additionally, partial and whole genome sequencing was attempted for any positive sample using RNA extracted directly from the diagnostic samples (*29*). For some swabs from longitudinally-sampled animals, the initial screening PCRs were conducted at the Wisconsin Veterinary Diagnostic Laboratory, followed by confirmatory testing at NVSL for any non-negative samples.

To explore other coronaviruses that may be present in the sampled animals, all respiratory swabs collected initially from each animal were subjected to a conventional RT-PCR to amplify a 668 bp-region within the RdRp gene that encodes the most conserved protein domain of α-, β-, γ-, and δ-coronaviruses (*30*). Amplicons were purified and submitted for bi-directional sequencing (Eton Biosciences, San Diego, CA) followed by manual editing and submission to Genbank (accession nos. MW263334-7).

### Viral Isolation

Specimens considered positive by qRT-PCR were subjected to virus isolation as described previously (*31*). Briefly, the samples were diluted 1:2 to 1:3 in minimum essential medium with Earle’s balanced salt solution (MEM-E). Vero (ATCC CCL-81) cells were inoculated with 1.5mL of diluted sample and adsorbed for 1 hour at 37°C. After adsorption, replacement medium was added and cells were incubated at 37°C for up to seven days. Cell cultures with no CPE were frozen, thawed and subjected to two blind passages, with inoculation of fresh cultures with the lysates as described above. Viral infection was confirmed through reduction of Ct values in the cell cultures with SARS-CoV-2-specific qRT-PCR using the CDC N1 and N2 primer and probe sets.

### Whole genome sequencing and phylogenetic analysis

Libraries for whole genome sequencing were generated using the Ion AmpliSeq Kit for Chef DL8 and Ion AmpliSeq SARS-CoV-2 Research Panel (Thermo Scientific, Waltham, MA). Libraries were sequenced using an Ion 520 chip on the Ion S5 system using the Ion 510™ & Ion 520™ & Ion 530™ Kit. Sequences were assembled using IRMA v. 0.6.7 (*32*) and visually verified using DNAStar SeqMan NGen v. 14. MAFFT (*33, 34*) was used to align FASTA files using a maximum number of iterative refinement number of 1,000. The alignment was used to output a phylogenetic tree using RAxML with GTRCAT model (*25*). The tree was rooted with the Wuhan seafood market reference genome (NC_0445512) and additional whole genome sequences from feline and canine sequences from across the world. Sequences from people from Texas were retrieved from NCBI and added to the analysis.

### Virus neutralization assay

All serum samples collected were assayed for SARS-CoV-2 neutralizing antibodies through virus neutralization (VN). For VN, 25 μL of two-fold serially diluted sera (for final dilutions with virus of 1:8 to 1:2048) were pre-incubated with 25 μL of 100 TCID_50_/ml of SARS-CoV-2 (2019-nCoV/USA-WA1/2020) in MEM-E containing 200 UI/mL penicillin, 200 μg/mL streptomycin, 75 μg/ml gentamicin sulfate and 6 μg/mL Amphotericin B for 60 min at 37°C with 5% CO_2_. Each serum sample was tested in duplicate in 96-well plates. At 1 hr post infection, 150 μL of Vero 76 cell suspension were added to the virus-serum mixtures. The neutralization titers were determined at 3 days post infection. The titer of a sample was recorded as the reciprocal of the highest serum dilution that provided at least 100% neutralization of the reference virus, as determined by visualization of CPE.

### National and International Reporting

The USDA case definition for a confirmed positive case of SARS-CoV-2 in animals includes PCR detection of both N1 and N2 at NVSL and sequence confirmation of virus either directly from the specimen or from virus isolate, or demonstration of SARS-CoV-2 neutralizing antibody (*35*). Confirmed positive cases were reported to CDC and USDA (*36*). USDA subsequently reported to the World Organisation for Animal Health (OIE) (*4*).

## Acknowledgments

We appreciate the participation of Texas pet owners. We thank Santos Navarrette, Robert Lampkin, and the Brazos County Health Department case investigators for their assistance in facilitating communication with COVID-19 cases. We thank Dr. Susan Rollo of Texas Animal Health Commission and Drs. Laura Robinson and Paul Gruenwald of Texas Department of State Health Services for assistance in reporting cases. Jennie Lamb and Jennifer Gauntt in CVMBS Communications provided project website development. Dr. Aliam Lim and Dan Christensen of the Wisconsin Veterinary Diagnostic Laboratory provided support in testing of selected re-samples from animals. We appreciate the assistance of Drs. Jessica Bourquin and Christine McFarland of TAMU Office of Biosafety while establishing field and lab protocols. The findings and conclusions in this report are those of the authors and do not necessarily represent the official position of the Centers for Disease Control and Prevention.

## Author contributions

S.A.H., A.P-C., and G.L.H. designed the study. S.A.H., I.B.Z., E.D., Y.A., and R.S.B.F. identified homes with animals to sample. S.A.H., I.B.Z., E.D., and L.D.A. sampled animals. C.M.R, W.T, M.T., M.L.K, M.J-M, and K.M. performed diagnostics. S.A.H., M.T., R.R.G., J.S., C.B.B., and G.L.H provided resources. All authors contributed to interpreting the results and editing the manuscript.

## Data and materials available

All data is available in the main text or the supplementary materials. Whole genome sequences of SARS-CoV-2 from animals obtained in this study are available at GenBank. Isolated SARS-CoV-2 from a cat is available from USDA upon request.

